# Cell shape, and not 2D migration, predicts ECM-driven 3D cell invasion in breast cancer

**DOI:** 10.1101/2019.12.31.892091

**Authors:** Janani P. Baskaran, Anna Weldy, Justinne Guarin, Gabrielle Munoz, Michael Kotlik, Nandita Subbiah, Andrew Wishart, Yifan Peng, Miles A. Miller, Lenore Cowen, Madeleine J. Oudin

## Abstract

Metastasis, the leading cause of death in cancer patients, requires the invasion of tumor cells through the stroma in response to migratory cues, such as those provided by the extracellular matrix (ECM). Recent advances in proteomics have led to the identification of hundreds of ECM proteins which are more abundant in tumors relative to healthy tissue. Our goal was to develop a pipeline to easily predict which of these ECM proteins is more likely to have an effect on cancer invasion and metastasis. We evaluated the effect of 4 ECM proteins upregulated in breast tumor tissue in multiple human breast cancer cell lines in 3 assays. We found there was no linear relationship between the 11 cell shape parameters we quantified when cells adhere to ECM proteins and 2D cell migration speed, persistence or 3D invasion. We then used classifiers and partial-least squares regression analysis to identify which metrics best predicted ECM-driven 2D migration and 3D invasion responses. ECM-driven 2D cell migration speed or persistence did not correlate with or predict 3D invasion in response to that same cue. However, cell adhesion, and in particular cell elongation and irregularity accurately predicted the magnitude of ECM-driven 2D migration and 3D invasion in all cell lines. Testing predictions revealed that our models are good at predicting the effect of novel ECM proteins within a given cell line, but that ECM responses are cell-line specific. Overall, our studies identify the cell morphological features that determine 3D invasion responses to individual ECM proteins. This platform will help provide insight into the functional role of ECM proteins abundant tumor tissue and help prioritize strategies for targeting tumor-ECM interactions to treat metastasis.

**Funding:** This work was supported by the National Institutes of Health [R00-CA207866-04 to M.J.O.]; Tufts University [Start-up funds from the School of Engineering to M.J.O.] and funds from NSF REU to A.W.

Conflict-of-interest: None.

**Insight Box:** Metastasis, the dissemination of tumor cells, is driven by the interaction of invading tumor cells with their local environment, in particular with the ECM, which provides structure and support to our tissues. This study presents an integrated approach to predict the effect of individual ECM proteins on 3D invasion and metastasis based on simple adhesion assays which quantify cell shape. Machine learning classification and partial-least squares regression models reveal that ECM-driven 2D cell migration metrics are not predictive of 3D invasion, and that cell shape of cells adhered to ECM can predict that protein’s effect on 3D invasion. These data provide a pipeline for predicting the effect of ECM proteins on breast cancer cell invasion and metastasis.

## Introduction

Metastasis, the dissemination of cells from the primary tumor to secondary organs in the body, is the leading cause of death in cancer. Metastasis involves the local invasion of tumor cells into the surrounding tissues, intravasation into the vasculature and lymphatics, and colonization of a distant site. All steps within tumor progression require cell migration – growth, invasion (1) and metastatic outgrowth (2). Understanding the mechanisms that drive cell migration in cancer is essential to identify strategies to treat cancers more effectively. Within tumors, several chemical and biophysical cues have been shown to promote local invasion (3). In particular, the extracellular matrix (ECM), which provides structure and support to our tissues, drives local invasion of tumor cells and metastasis, as well as colonization of secondary sites. For example, the glycoprotein Fibronectin, which is produced by both tumor and stromal compartments in breast tumors (4), can drive directional migration of breast cancer cells to drive metastasis (5). The optimization of protocols to characterize the ECM of tumors has led to the identification of multiple ECM proteins abundant in tumor tissue that may be involved in promoting metastatic phenotypes (4, 6). The present study aims to develop a pipeline to easily assess which of these ECM proteins, alone or in combination, are more likely to affect invasion and metastasis, and are therefore better targets as biomarkers or for drug development.

Breast cancer cells sense ECM cues within their environment via cell surface receptors and the extension of actin-rich protrusions such as lamellipodia and filopodia. The activation of downstream signaling pathways and the formation of focal adhesions promotes cytoskeletal dynamics which help the cell propel itself forward, eventually retracting its tail via disassembly of focal adhesions (7–10). Cell-ECM interactions and their impact on cell behavior can be studied in different contexts. Cell responses to ECM cues have been measured as alterations in cell shape following adhesion to a substrate, 2D migration on a substrate, and 3D invasion into a matrix containing the ECM substrate. However, we still do not understand the relationship between adhesion to, 2D migration on, and 3D invasion in a given ECM substrate. Therefore, there is a critical need to create a predictive model to use cell morphology to predict cell invasion responses to ECM cues.

Existing models that predict cell migration have focused on cell morphology or signaling pathways and mostly focused on a single cue. First, cell morphology or shape is commonly used to characterize cellular phenotypes, because it can be easily visualized and quantified using traditional immunostaining and basic microscopy. Epithelial keratocytes from fish skin have been used to generate various models due to their characteristic and homogeneous fan-like shape. Various models have been published linking cell shape and geometry with cell migration and speed (11, 12). This has been more challenging for cancer cells given their more complex and heterogeneous cell morphologies. There have been efforts to identify signaling pathways that regulate cell morphology. One study linked breast cancer cell morphology *in vitro* in 3D Matrigel with gene expression to identify dominant genes that are predictive of morphological features (13). Quantitative morphological profiling has also been used to evaluate the role of individual genes in regulating cell shape using genetic screens in drosophila cells, leading to the identification of signaling networks that regulate cell protrusion and adhesion (14). However, these studies all focus on a single ECM cue, and there are currently no studies predicting mesenchymal 3D cell movement in response to ECM cues.

The goal of this study is to understand how ECM cues in the tumor microenvironment promote invasion and metastasis of cancer cells from the primary tumor. Using classifier-based and partial-least square regression models trained on cell morphology, 2D migration, and 3D invasion data, we find that cell shape in response to a particular ECM protein can predict the ability of a cell line to invade through that ECM protein in 3D.

## Methods

### ECM substrates

Reagents were purchased from Fisher Scientific (Hampton, NH) or SIGMA (St. Louis, MO) unless otherwise specified. We used the following ECM proteins: Collagen I protein (CB-40236; Fisher Scientific, Hampton, NH), Fibronectin protein (F1141; SIGMA, St. Louis, MO), Tenascin C (R&D systems, 3358TC050), Collagen IV protein (Abcam, ab7536) and Matrigel (growth-factor reduced, Corning, CB-40230C).

### Cell culture

MDA-MB-231, MDA-MB-468 and BT549 cells were obtained from ATCC (Manassas, VA). MDA-MB-231 and MDA-MB-468 cells were cultured in DMEM with 10% serum and Pen-Strep Glutamine and BT549 were grown in RPMI +10%PBS +Insulin (1μg/mL). Cells were checked every 2 months for the presence of mycoplasma by a PCR based method using a Universal Mycoplasma Detection Kit (30-1012K; ATCC, Manassas, VA). Only mycoplasma negative cells were used in this study. Cell lines were made to stably express GFP by lentiviral transduction and labeled as 231-GFP or 468-GFP.

### Adhesion assay

Plastic-bottomed dishes (Thermo Fisher Nunc, 96 Well Optical-Bottom Plates, 165305) were coated with 20μg/ml ECM protein for 1hr at 37°C and then washed with PBS. Cells were trypsinized, resuspended in media and plated on the coated plates at 4,000 cells/per well. After 2hrs, cells were then fixed for 15 min in 4% paraformaldehyde, then permeabilized with 0.2% TritonX-100, blocked with 3% BSA and incubated with primary antibodies overnight at 4°C. Cells were DAPI (D1306; Thermo Fisher Scientific, Waltham, MA) and Phalloidin (A12390; Thermo Fisher Scientific, Waltham, MA) stained along with incubation with fluorescently labeled secondary antibodies at room temperature for 2 hours. Imaging was performed using a Keyence BZ-X710 microscope (Keyence, Elmwood park, NJ) and CellProfiler v3.1.8 was used for imaging analysis using a custom pipeline(15). Cells were first identified from the nucleus, and the outline of each cell was determined from the cytoplasm staining. Cells at the edge of an image were discarded. 2D Adhesion was quantified by 11 parameters: area/cell (number of sq um in the cell cytoplasm), aspect ratio (the ratio of the major axis length and the minor axis length of the cell), compactness (mean squared distance of the cell cytoplasm from the centroid divided by the area, where a filled circle has a value of 1, and an irregular shape has a value greater than 1), eccentricity (ratio of the distance between the foci of the ellipse and its major axis length, where a perfect circle has a value of 0, and more elongated cells have a value of 1), extent (proportion of pixels in the bounding box that are also in the cell cytoplasm, where larger values indicate more spread out cell cytoplasm), form factor (calculated as 4*π*Area/Perimeter^2^, where a perfect circle has a value of 1), max. and min feret diameter (minimum and maximum distance between two parallel lines that are tangent to the cell cytoplasm edge), mean radius, perimeter, and solidity (proportion of pixels that are in the convex hull that are also in the cell cytoplasm, where a perfect circle has a value of 0). Data are the result of 3 independent experiments with 3 technical replicates per experiment.

### 2D migration assay

For 2D migration, 12-well glass-bottomed dishes (MatTek, Ashland, MA) were coated with 20μg/ml ECM protein for 1hr at 37°C. ECM was washed off with PBS, and cells were plated at 7,500 cells/well on and allowed to adhere. After 1hr, the plate was placed on the microscope and cells were imaged overnight with images acquired every 10 min for 16 hours in an environmentally controlled chamber within the Keyence BZ-X710 microscope (Keyence, Elmwood park, NJ). Cells were then tracked using VW-9000 Video Editing/Analysis Software (Keyence, Elmwood park, NJ) and both cell speed and persistence calculated using a custom MATLAB script vR2018a (MathWorks, Natick, MA). 2D migration was quantified by 2 parameters: cell migration speed and persistence. Data are the result of 3 independent experiments with 6 fields of view per experiment and an average of 6 cells tracked per field of view.

### Spheroid invasion and migration assay

Cells were seeded in low-attachment plates in media (Corning™ 96 Well Ultra-Low Attachment Treated Spheroid Microplate, 12-456-721), followed by centrifugation at 3,000 rpm for 3 mins to form spheroids. Spheroids were grown for 3 days after which ECM was added to each well, which included (depending on the condition) Collagen I protein to a 1mg/ml concentration, ECM protein of interest at 20μg/ml, 10mM NaOH, 7.5% 10x DMEM and 50% 1x DMEM. The spheroids in ECM were then spun down at 3,000 rpm for 3 mins, and the ECM gel left to polymerize for 1hr at 37°C. After this, a further 50μl of media added to each well. Following another 5 days of growth, spheroids were imaged as a Z-stack using a Keyence BZ-X710 microscope (Keyence, Elmwood park, NJ) and Z-projection images analyzed using a Hybrid Cell Count feature within the BZ-X Analyzer software v1.3.1.1 (Keyence, Elmwood park, NJ). 3D invasion was quantified by 1 parameter: increase in surface area on Day 5 of ECM relative to Day 1. Data are the result of 3 independent experiments with 6 technical replicates per experiment.

### Clustering Analyses

SPRING plots were constructed from single cell adhesion quantification using the methods described in Weinreb *et al.* (16). For large scale quantification of cell adhesion on different ECM substrates, each profile was averaged and mean centered. The ECM factors and adhesion metrics were clustered by rank correlation and mean linkage, using the seaborn package for Python. Cell adhesion shape metrics were compared by calculating the Spearman correlation between each pair of metrics.

### Machine Learning Classification

ECM-driven effects on 2D migration and 3D invasion were classified as either low or high (see Fig 5). For classification of 2D migration in MDA-MB-231 and MDA-MB-468 cells, ECM substrates that caused a mean cell migration speed of above 0.5μm/min were classified as high. For classification of 3D invasion in MDA-MB-231 cells, ECM substrates that caused a mean fold change in spheroid area of above 10 were classified as high. For classification of 3D invasion in MDA-MB-468 cells, ECM substrates that caused a mean fold change in spheroid area of above 8 were classified as high. Machine learning algorithms, Support Vector Machine (SVM), Linear Support Vector Machine (Linear SVM), Random Forest (RF), and Logistic Regression (LR), were trained and validated using the classified data, and model parameters optimized in cross validation using a grid search. AUROC area under the curve was used to assess the accuracy of the classifiers. The optimized models were tested using a new unknown ECM protein. All machine learning classification was performed using Python.

### Principal Component Analysis and Partial Least-Squares Regression

Principal component analysis and partial least-squares regression were performed as described previously (17). Model predictions were made by introducing a new condition (ECM protein or cell line). Model fitness was calculated using R^2^, RMSE, and percent error, previously described (18). VIP scores were calculated from reference (19). Adhesion parameters with VIP scores above 1 were considered as important cell adhesion parameters for prediction. All data was scaled to nondimensionalize the different metrics. PCA was performed using Python, and PLS model was implemented using Matlab.

### Statistical analysis

GraphPad Prism v7.04 was used for generation of graphs and statistical analysis. All statistical comparisons were done between no ECM condition and each ECM condition individually. A one-way ANOVA was used with a corrected p-value of ≤ 0.05 considered significant.

## Results

### ECM impacts breast cancer cell adhesion

We chose to build our model using 4 ECM proteins known to be abundant in breast tumors: Collagen I, Collagen IV, Fibronectin, and Tenascin C. Indeed, these components were identified in xenograft 4T1 breast tumors (6) and in highly metastatic LM2 tumors (4). Collagen I and the glycoprotein Fibronectin are two of the most abundant ECM proteins in mammary tumors, and both are known to contribute to breast cancer invasion and metastasis (5, 20). Another glycoprotein, Tenascin C, has also been shown to contribute to breast cancer metastasis (21). Collagen IV is a major component of the basement membrane, which breast cancer cells must break down to invade surrounding tissues (22, 23). To investigate the effects of these ECM proteins, we used two human triple-negative breast cancer cell lines, MDA-MB-231 and MDA-MB-468. MDA-MB-231 is mesenchymal, with high metastatic potential in mouse models, while MDA-MB-468 is epithelial, with lower metastatic potential (24, 25).

First, we performed an adhesion assay, which has been commonly used to study cell-ECM interactions, where cells are plated on a 2D ECM-coated surface and left to adhere for 2hrs. The cells are then fixed and immunostained to assess cell shape. We focused our efforts on the actin cytoskeleton, given that cell shape is associated with adhesion and cell migration (Fig 1A). We quantified 11 cell shape parameters via Cell Profiler, and established effects of all 4 ECM proteins on these parameters. Collagen I, Fibronectin and Collagen IV led to increased cell area, eccentricity, which characterizes cell elongation, and compactness, which quantifies cell shape irregularity. Tenascin C decreased cell area, eccentricity and compactness (Fig 1B-D). While MDA-MB-468 cells had a smaller cell area on average, the effect of individual ECM proteins had similar relative effects on cell morphology (Fig 1E-H). These data comprehensively characterize the effect of 4 ECM proteins upregulated in breast tumor tissue on breast cancer cell shape.

**Figure 1:**
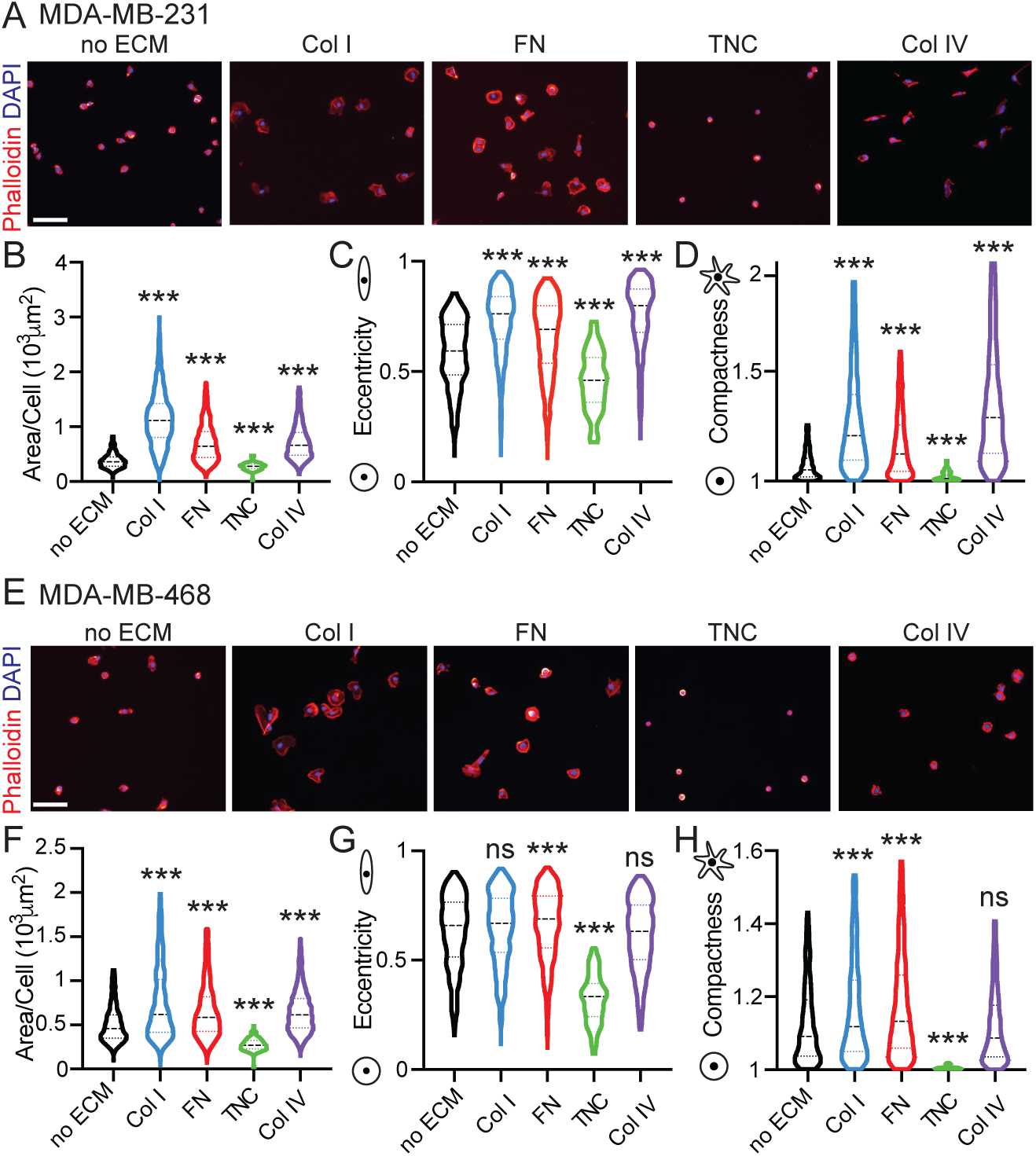
ECM proteins upregulated in breast tumor tissue have distinct cell line-specific effects on tumor cell adhesion. A) Representative images of MDA-MB-231 cells plated on plastic, Collagen I, Fibronectin, Tenascin C or Collagen IV for 2hrs, fixed and stained with Phalloidin (red) and DAPI (blue). Scale bar is 100μm. Quantification of cell shape features using Cell Profiler to evaluate effects on MDA-MB-231: B) area/cell (10^3^μm^2^), C) eccentricity and D) compactness. Results show entire distribution and significance by one-way ANOVA, ***p<0.005. E) Representative images of MDA-MB-468 cells plated on plastic, Collagen I, Fibronectin, Tenascin C or Collagen IV for 2hrs, fixed and stained with Phalloidin (red) and DAPI (blue). Scale bar is 100μm. Quantification of cell shape features using Cell Profiler to evaluate effects on MDA-MB-468: F) area/cell (10^3^μm^2^), G) eccentricity and H) compactness. Results show entire distribution and significance by one-way ANOVA, ***p<0.005, ns is not significant.

To better visualize the effects of ECM proteins on shape parameters, we used SPRING, a pipeline for data filtering, normalization and visualization using force-directed layouts of k-nearest neighbor algorithms (Fig 2A). SPRING has been shown to reveal more detailed biological relationships than existing approaches, with plots being more reproducible than those of stochastic visualization methods such as tSNE (16). Individual ECMs were mapped onto the individual cells on the SPRING plot (Fig 2B). For the MDA-MB-231 cells, as was seen in the clustering, cells on Tenascin C and no ECM cluster together. Interestingly, cells on Collagen I are seen as very distinct from cells on no ECM, with little overlap. Cells on Collagen IV are also distinct from cells on no ECM, but are also separate from the cells on Collagen I. Finally, the cells on Fibronectin are homogeneously distributed throughout the cluster.

**Figure 2:**
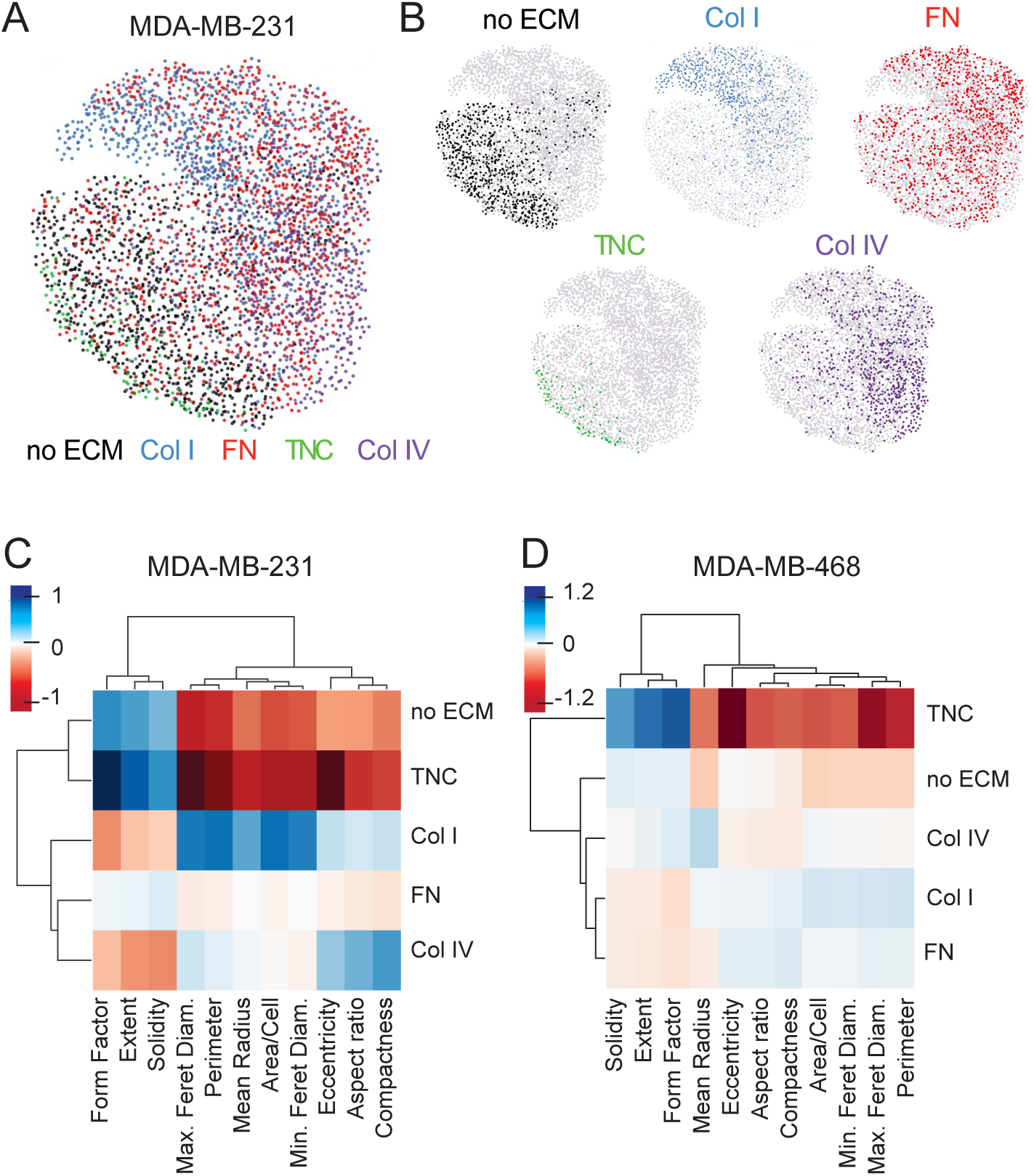
Clustering of adhesion parameters reveals ECM-specific effects on cell shape. A) Visualization of continuum of cell adhesion of MDA-MB-231 cells on different ECM substrates based on all 11 cell shape parameters with SPRING plots. B) Plots showing localization of ECM factor-dependent cell adhesion shape on the combined SPRING plot. Mean centered cell adhesion of MDA-MB-231 (C) and MDA-MB-468 (D) cells plated on plastic, Collagen I, Fibronectin, Tenascin C or Collagen IV for 2hrs. Each cell adhesion parameter and ECM factor is clustered by rank correlation and mean linkage.

Initial analysis of the entire dataset by unsupervised clustering demonstrated that ECM-driven effects on shape parameters cluster into 3 main groups: one with compactness, aspect ratio and eccentricity, which quantify how elongated is a cell is, a second group with solidity, form factor and extent, which describe how irregular the shape of the cell is or how protrusive a cell is; and a third group with the ferret diameters, radius, perimeter and cell area, which quantify how large a cell is. Both cell lines clustered in these groups (Fig S1A,B). While the effect of the different ECM proteins on these parameters clustered differently in each cell line, it is clear that Tenascin C shape quantifications are more similar to the no ECM shapes, while Fibronectin and Collagen IV shape characteristics tend to cluster with each other (Fig 2C,D). Overall, these clustering methods demonstrate ECM proteins have distinct effects on cell shape.

### ECM-driven 2D migration does not correlate with cell shape

We then investigated the effect of these same ECM cues on 2D cell migration, by evaluating cell speed and persistence. Cell speed measures how fast a cell is moving over a given distance, while persistence, the Euclidian distance between start and finish over the total distance traveled, informs whether the cell is moving in a straight line (closer to 1) or taking a more winding path. In MDA-MB-231 cells, we find that Collagen I and Collagen IV increase both cell migration speed and persistence (Fig 3A-C). Fibronectin has no effect on cell speed or persistence; Tenascin C decreases cell migration speed while increasing persistence. We find that for all ECM conditions there was no significant correlation between cell migration speed and persistence (Fig 3D), or between cell area and cell migration speed (Fig 3E). Similar results were obtained with the MDA-MB-468 cell line, where Collagen I and Collagen IV increased cell migration and persistence, with Tenascin C reduced cell migration and persistence (Fig 3F-H). There was also no significant correlation between cell migration speed and either persistence or cell area (Fig 3I-J). Overall, these findings suggest that ECM-driven effects on cell speed and persistence are distinct, and that effects on cell shape may not correlate with effects on cell migration speed.

**Figure 3:**
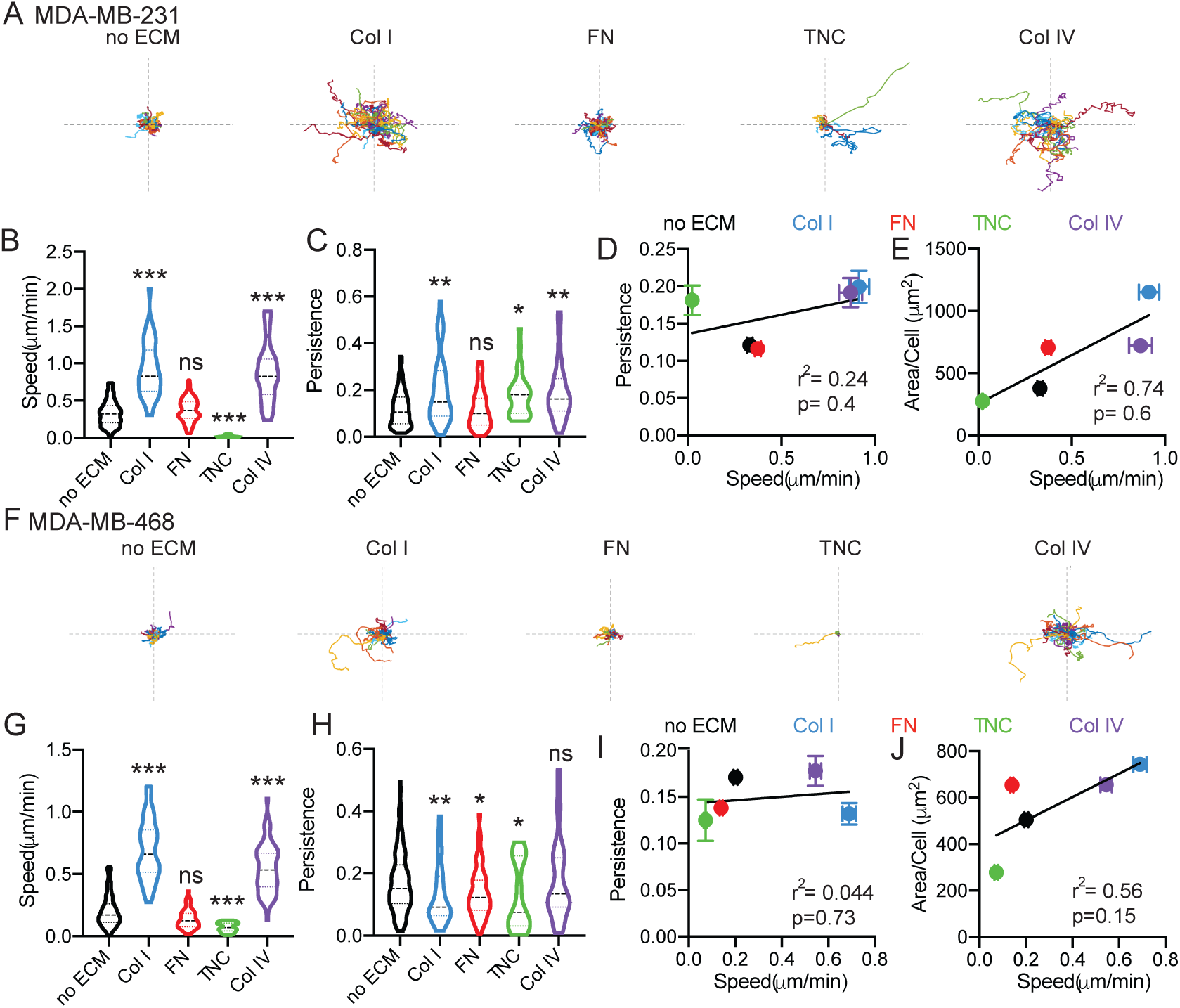
ECM-driven effects on 2D cell migration speed do not correlate with effects on persistence. A) Representative roseplots for MDA-MB-231 cells plated on glass, Collagen I, Fibronectin, Tenascin C or Collagen IV for 16hrs and imaged every 10 mins. Each line represents an individual cell. Axis length is 500μm. Quantification of cell migration speed (μm/min) (B) and persistence (C). Correlation between 2D persistence and 2D cell migration speed (D) and between cell area (μm^2^) and 2D cell migration speed (E). F) Representative roseplots for MDA-MB-468 cells plated on glass, Collagen I, Fibronectin, Tenascin C or Collagen IV for 16hrs and imaged every 10 mins. Each line represents an individual cell. Axis length is 500μm. Quantification of cell migration speed (μm/min) (G) and persistence (H). Correlation between 2D persistence and 2D cell migration speed (I) and between cell area and 2D cell migration speed (J). Results show entire distribution and significance by one-way ANOVA, * p<0.05, ** p<0.01, ***p<0.005, ns is not significant. Data pooled from at least 3 independent experiments, between 30-130 individual cells analyzed per condition.

### ECM-driven 3D invasion does not correlate with 2D migration

It is well established that 3D invasion is a more physiologically relevant model of *in vivo* cell migration, therefore we quantified the effect of the individual ECM proteins on 3D invasion in Collagen I gels. We used a spheroid model of 3D invasion, which constitute microtumors recapitulating various clinically important characteristics like hypoxia, nutrient- and pH gradients and deposition of ECM. All spheroid gels contain Collagen I as matrix to support spheroid formation, given that it is the most abundant ECM component of breast tissue, and that all ECM proteins present in tumors would be in the presence of Collagen I. We find that all 4 ECM proteins drive a significant increase in invasion relative to Collagen I only in MDA-MB-231 cells (Fig 4A,B). However, we do not find a significant correlation between the effect of these proteins on 2D migration and 3D invasion (Fig 4C). Similarly, the 4 ECM proteins also increase invasion of MDA-MB-468 cells (Fig 4D,E), although these cells migrate more individually than the MDA-MB-231 cells. There is also no significant correlation between ECM effects on 2D cell migration and 3D invasion (Fig 4F).

**Figure 4:**
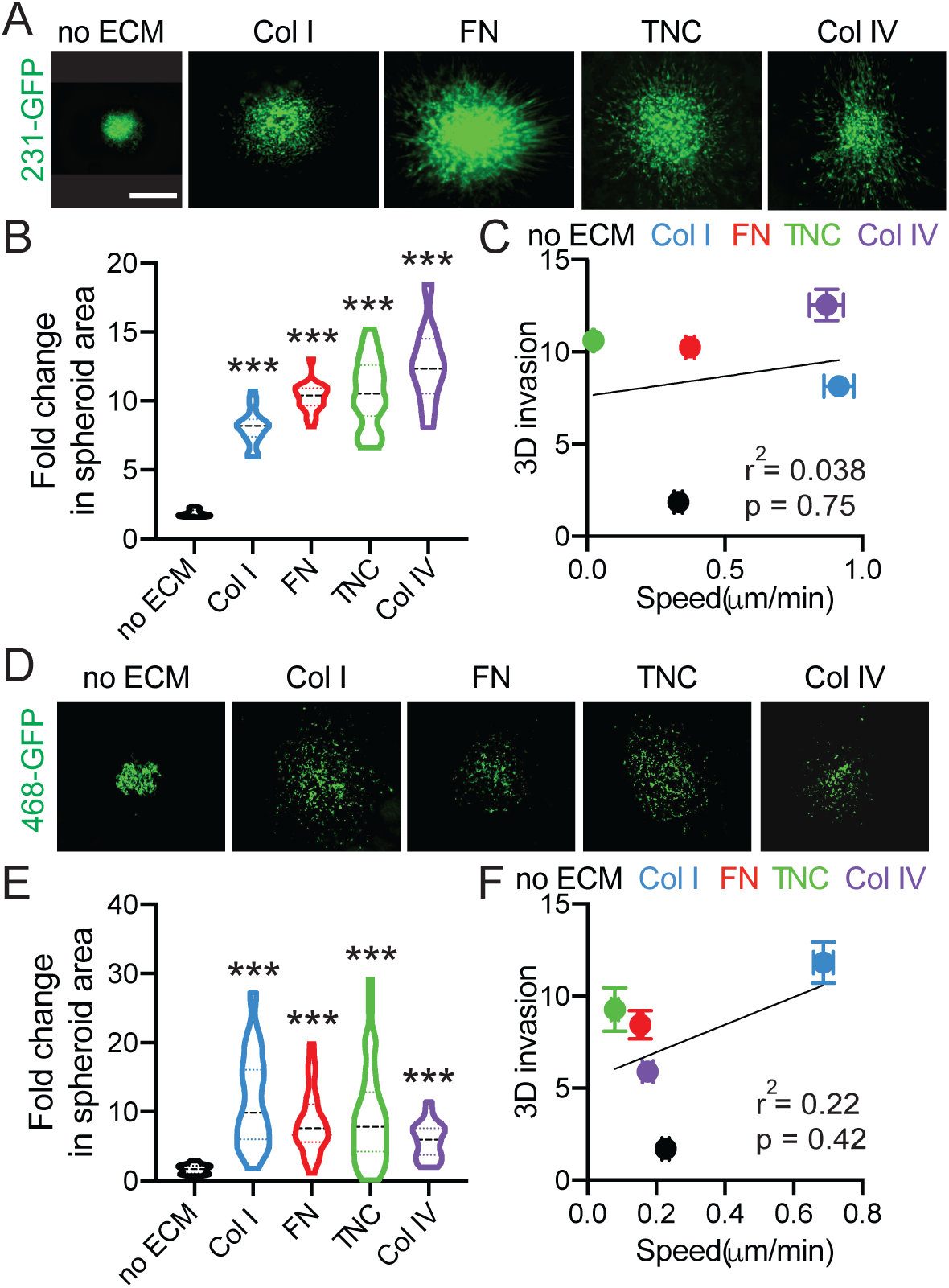
ECM-driven 3D invasion does not correlate with effects on 2D cell migration. A) Representative images of spheroids made from 231-GFP cells embedded in media, Collagen I, Fibronectin, Tenascin C or Collagen IV gels for 5 days. Scale bar is 200μm. B) Quantification of fold change in 231-GFP spheroid area on day 5 relative to day 1. Data pooled from at least 5 biological replicates, with technical triplicates. *** p<0.001 by one-way ANOVA and Dunn’s multiple comparison test. C) Correlation between mean fold change in spheroid area and 2D cell migration speed for 231-GFP cells embedded in media, Collagen I, Fibronectin, Tenascin C and Collagen IV. D) Representative images of spheroids made from 468-GFP cells embedded in media, Collagen I, Fibronectin, Tenascin C or Collagen IV gels for 5 days. Scale bar is 200μm. E) Quantification of fold change in 468-GFP spheroid area on day 5 relative to day 1. Data pooled from at least 4 biological replicates, with technical triplicates. *** p<0.001 by one-way ANOVA and Dunn’s multiple comparison test. F) Correlation between mean fold change in spheroid area and 2D cell migration speed for 468-GFP cells embedded in media, Collagen I, Fibronectin, Tenascin C and Collagen IV.

### Generation of classifier-based model suggests that cell adhesion can be used to categorize 2D migration and 3D invasion

To dissect the relationship between ECM-driven cell adhesion, 2D migration, and 3D invasion and develop methods to predict ECM-driven effects on breast cancer cells, we first used machine learning classifier models. Our goal was to evaluate the ability of different learning algorithms to predict 2D migration based on cell adhesion parameters and 3D invasion based on either adhesion parameters or 2D migration. We assigned each ECM protein as either as ‘low’ or ‘high’, based on its ability to induce 2D cell migration or 3D invasion. Based on the results in Figure 3, 2D cell migration speed of MDA-MB-231 cells on no ECM, Fibronectin, and Tenascin C was classified as low (Fig 5A). Based on the results from Figure 4, 3D invasion of MDA-MB-231 cells embedded in no ECM and Collagen I was classified as low, while 3D invasion in Fibronectin, Tenascin C and Collagen IV was classified as high (Fig 5C,E). We used 4 different commonly used machine learning algorithms to classify the data. First, we used Logistic Regression (LR), a statistical method for analyzing a dataset in which there are one or more independent variables that determine an outcome through fitting a logistic function; this outcome is measured as one of 2 outcomes. We used a support-vector machine (SVM) algorithm, which is a classification method that samples hyperplanes to separate between two classes. We used a linear SVM and SVM in our analysis, where the linear SVM assumes that the data set distribution can be linearly divided. Finally, we used Random Forest (RF), which creates decision trees on randomly selected data samples. This algorithm takes averages of predictions from each tree and uses them to further refine the model to improve results while also minimizing over-fitting. The ability of an algorithm to accurately predict whether an ECM protein has a low or high effect is assessed via the Area Under the Curve Receiver Operating Characteristic (AUROC) score, a performance measurement for classification problems.

First, we assessed the predictive relationship between cell adhesion and 2D cell migration in response to different ECM proteins. Interestingly, using all 11 cell shape parameters were able to predict 2D migration, with an AUROC score higher than 0.75. (Fig 5B). We then tested whether any of the groups of cell features identified in Figure 2, cell size, irregularity and elongation could independently predict 2D migration. We found that the cell size parameters (area/cell, perimeter, mean radius, min and max feret diameter) could also accurately predict 2D migration speed with AUROC scores above 0.75, while cell elongation and irregularity could not. While all 4 algorithms performed similarly, the SVM algorithm performed the best. We found similar results with the MDA-MB-468 cells, where for 2D cell migration, we classified no ECM, Tenascin C and Fibronectin as low, and Collagen I and Collagen IV as high (Fig S4A). All 11 parameters were able to accurately predict 2D cell migration with an AUROC over 0.75 (Fig S4B). These data demonstrate that cell shape of cells adhered to a particular ECM protein is a reliable metric for predicting how this protein will impact 2D migration speed.

Next, we determined the predictive relationship between 2D cell migration and 3D invasion in response to different ECM proteins. For both MDA-MB-231 and MDA-MB-468 cells, the AUROC scores for the optimized classifier models are insignificant (around 0.5) suggesting that the classifiers were operating with a similar accuracy to that of a random classification assignment (Fig 5D, Fig S4D). Interestingly, for both cell lines, models using cell migration speed alone tended to have higher AUROC scores than those using persistence, suggesting that of the two parameters, speed was better at predicting 3D invasion. This result further supports initial regression analysis that suggests there is no clear predictive relationship or correlation between 2D migration and 3D invasion.

Finally, we assessed the predictive relationship between cell adhesion and 3D cell invasion in response to different ECM proteins. For both MDA-MB-231 and MDA-MB-468 models, using all shape parameters yielded the highest AUROC scores (Fig 5F, Fig S4F). Models constructed with cell size parameters, cell irregularity parameters, and cell elongation parameters did not have significant AUROC scores. For the binary classification of MDA-MB-468 cells, the most promising result was an AUROC score of .775 when using all parameters and the SVM classifier. It is notable that the results from classifying with all parameters were all higher than the AUROC scores obtained from just relying on parameters associated with cell size. This demonstrates that cell adhesion can classify 3D cell invasion more accurately than 2D cell migration.

**Figure 5:**
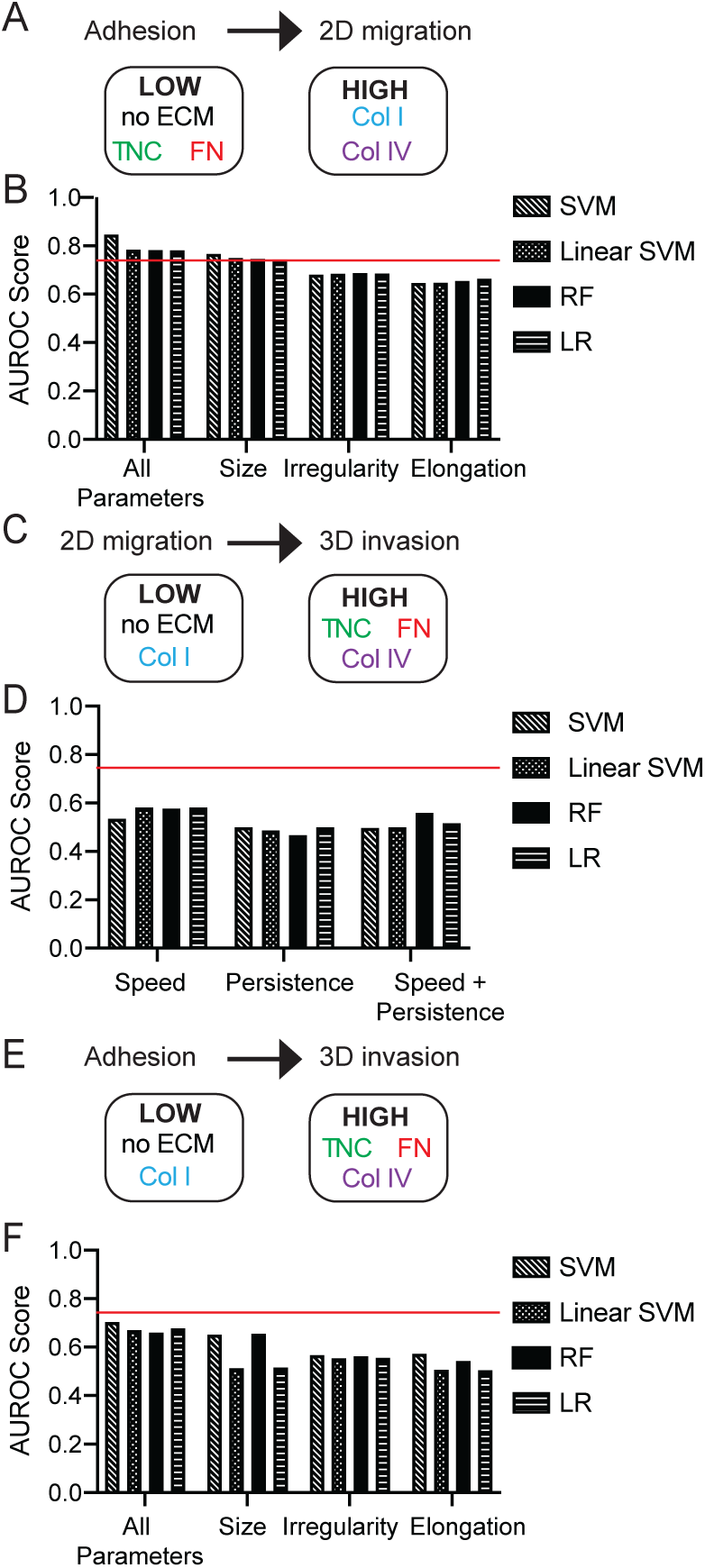
Adhesion classifies ECM-driven 2D migration and 3D invasion. A) Cell adhesion to predict binary classification of 2D cell migration speed of MDA-MB-231 cells on plastic, Collagen I, Fibronectin, Tenascin C, or Collagen IV. B) AUROC scores of binary classifier models (A) using all 11 cell shape parameters, cell size parameters (area/cell, perimeter, mean radius, min feret diameter, max feret diameter), cell irregularity parameters (solidity, extent, form factor), and cell elongation parameters (eccentricity, aspect ratio, compactness). C) 2D cell migration to predict binary classification of mean fold change of spheroid area of 231-GFP cells embedded in media, Collagen I, Fibronectin, Tenascin C, or Collagen IV. D) AUROC scores of binary classifier models (C) using 2D cell migration (cell migration speed and persistence). E) Cell adhesion to predict binary classification of mean fold change of spheroid area of 231-GFP cells embedded in media, Collagen I, Fibronectin, Tenascin C, or Collagen IV. F) AUROC scores of binary classifier models (E) using all 11 cell shape parameters, cell size parameters, cell irregularity parameters, and cell elongation parameters.

### PLS models suggest that cell adhesion accurately predicts 3D invasion

We used data-driven modeling to more precisely determine the relationship between ECM-driven cell adhesion, 2D migration and 3D invasion. First, we used principal components analysis (PCA) to reduce the dimensionality of the cell adhesion data set. The PCA creates a new set of principal components which maximize the covariance captured between the parameters {Janes, 2006 #416}. Using two principal components, over 80% of the variation in cell adhesion is described, and the distinct effects of each ECM substrate can be identified (Fig S5A-D). The PCA shows similar ECM-specific distributions of cell adhesion seen in the SPRING plots, indicating that even with reducing the data dimensionality the important trends are captured.

Next, we used a partial least-squares regression (PLSR) to identify covariation between cell adhesion and 2D migration and 3D invasion. The PLS model reduces the data to a set of principal components to optimally describe the proposed relationship between the input, cell adhesion, and the outputs, 2D migration and 3D invasion {Janes, 2006 #416}. We constructed the PLS model with MDA-MB-231 cells only (Fig 6), MDA-MB-468 cells only (Fig S6), and a combination of both cell lines (Fig S7). The scores plot of principal component one (PC1) and PC2 describes how strongly each ECM factor projects on each principal component (Fig 6A). For example in MDA-MB-231 cells, Collagen I and Collagen IV project negatively on PC1, whereas no ECM, Fibronectin, and Tenascin project positively on PC1. Therefore, using both PC1 and PC2, we can distinguish the variation between the effects of different ECM substrates. For the combined model, both PC1 and PC2 are required to describe variation between different cell lines and ECM substrates (Fig S7A). For no ECM, FN, and TNC, both cell lines project similarly onto the principal components. However, for Collagen I and Collagen IV, the cell lines project differently onto the principal components. To understand the effects of cell adhesion parameters in the model, we projected the loading vectors, which describes how strongly each parameter projects onto each principal component (Fig 6B). We find that cell size, irregularity and elongation project in distinct clusters, where cell irregularity projects positively on PC1 and cell size and elongation project negatively. To evaluate model fitness, we calculated the R^2^ to measure the percent of variance captured by the model and root mean square error (RMSE) to measure the deviation between the data and model prediction (Fig 6C). We determined the ideal number of principal components to use such that the RMSE is minimized and R^2^ is maximized, without overfitting. For the individual cell line models, we used 2 principal components, and for the combined cell line model we used 6 principal components.

**Figure 6:**
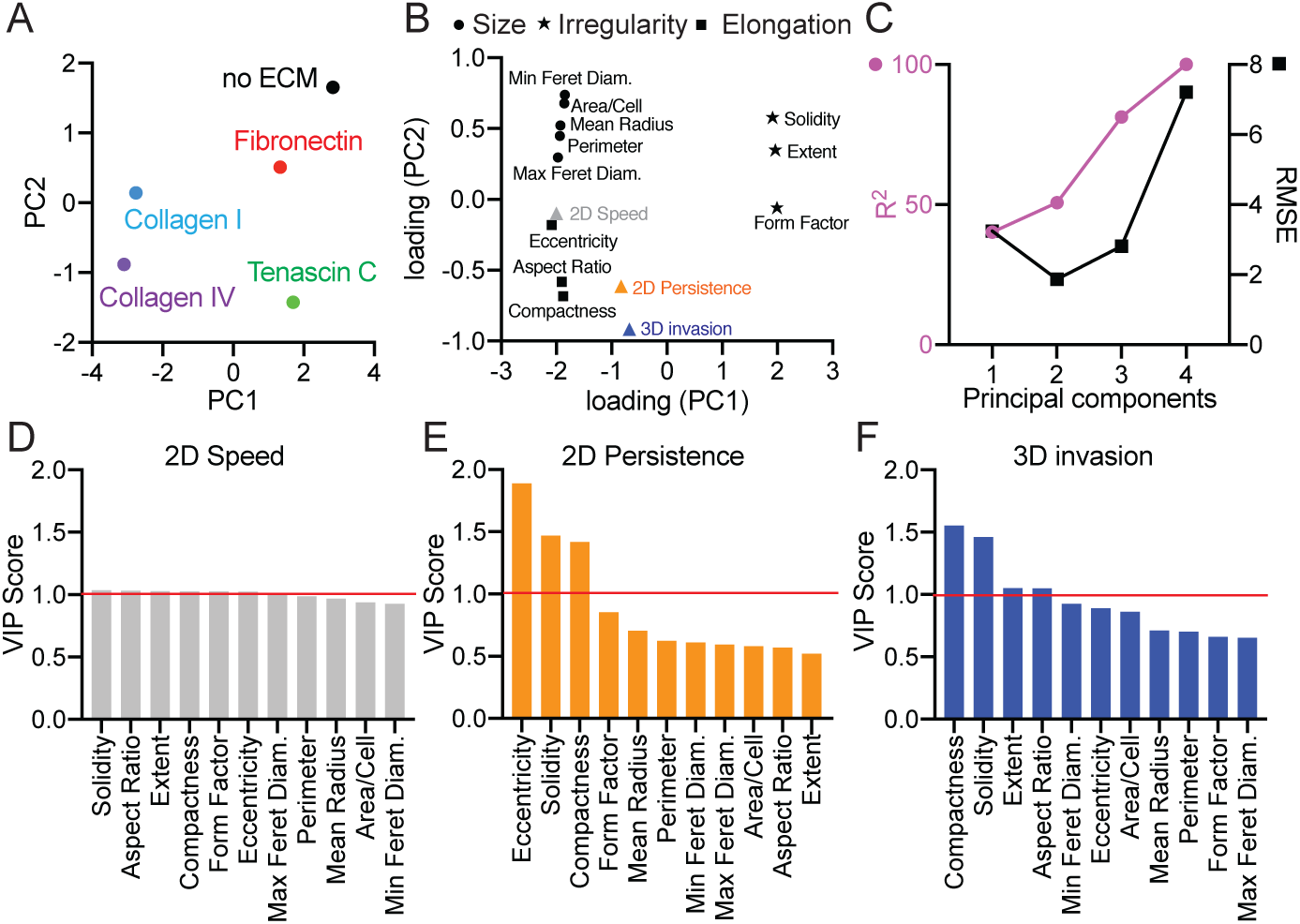
A partial least squares regression model constructed to predict ECM-driven 2D migration and 3D invasion from cell adhesion. A) Scores plot for PLS model with MDA-MB-231 cells. Principal components reflect covariation between adhesion, 2D migration and 3D invasion for each cell line on each ECM protein. B) PLS loading plot for 11 cell adhesion shape parameters and 3 cell responses. C) R^2^ and root mean square error (RMSE) for the PLS model built with increasing numbers of principal components. Ranked VIP scores for predicting 2D cell migration speed (D), 2D persistence (E), and 3D invasion (F). VIP score >1 indicates important cell shape parameters to predict cell invasion response.

We then identified how different cell adhesion parameters contribute to prediction of 2D speed, 2D persistence, and 3D invasion using the variance importance parameter (VIP) score for each cell adhesion parameter (Fig 6D-F). The VIP score reports the amount of variation in 2D speed, 2D persistence and 3D invasion that is explained by each adhesion parameter. We find that in the MDA-MB-231 model, all the cell adhesion parameters rank similarly for predicting 2D speed, indicating that all parameters are important for prediction (Fig 6D). However, in the MDA-MB-468 model, we find that mean radius, cell area, and min feret diameter, which are measures of cell size, are important for predicting 2D speed (Fig S6D). Similar to the MDA-MB-231 model, we find that in the combined cell line model, all parameters rank similarly for predicting 2D speed (Fig S7D). Interestingly, in the MDA-MB-231 and combined cell lines models, measures of cell irregularity and elongation are important for predicting 2D persistence and 3D invasion. In the MDA-MB-231 model, eccentricity, solidity, and compactness are important for predicting 2D persistence, and compactness, solidity, and extent are important for predicting 3D invasion (Fig 6E,F). In the combined cell line model, compactness, form factor, and solidity are important for predicting 2D persistence, and eccentricity, compactness, and solidity are important for predicting 3D invasion (Fig S7E,F). In the MDA-MB-468 model, we find that measures of cell size and elongation are important for predicting 2D persistence and 3D invasion (Fig S6E,F). Mean radius and eccentricity are important for predicting 2D persistence, and eccentricity, aspect ratio, cell area, and min feret diameter are important for predicting 3D invasion. Overall, these models demonstrate the importance of individual shape parameters in predicting ECM-driven migration responses in 2D and 3D.

### Models can be used to accurately predict 2D and 3D ECM-driven responses

We first tested the ability of our data-driven models to predict the effects of a new ECM protein, within the same cell line. We chose Matrigel, isolated from the Engelbreth-Holm-Swarm (EHS) mouse sarcoma, which is rich in basement membrane components laminin, collagen IV and heparan sulfate proteoglycans. We measured cell adhesion, 2D migration and 3D invasion of MDA-MB-231 and MDA-MB-468 cells in response to Matrigel (Fig 7A-B). First, we used the classifier models with all parameters to predict the cell responses to Matrigel, since using either all cell adhesion parameters or all cell migration parameters had the best AUROC scores (Fig 5). We find that the models accurately classify 2D migration and 3D invasion of cells on Matrigel from cell adhesion (Fig 7C, E). We also find that 2D migration accurately classifies 3D invasion (Fig 7D). However, the AUROC scores for the classifier models using 2D migration to predict 3D invasion were low, indicating that although the prediction of Matrigel was accurate, the model has low confidence in that classification. We then used the PLS models to quantitatively predict cell responses to Matrigel using all cell adhesion parameters (Fig 7F,G). In MDA-MB-231 cells, we find that the model predicts the effects of Matrigel accurately, as demonstrated by the low percent errors (Fig 7F). Interestingly, we find that only 2D persistence and 3D invasion, but not 2D speed, are well predicted in the MDA-MB-468 model (Fig 7G). Prediction of Matrigel from the combined model has a larger error, indicating cell line specificity (Fig S8G). Additionally, using both cell adhesion and 2D migration to predict 3D invasion increased error (Fig S8H).

**Figure 7:**
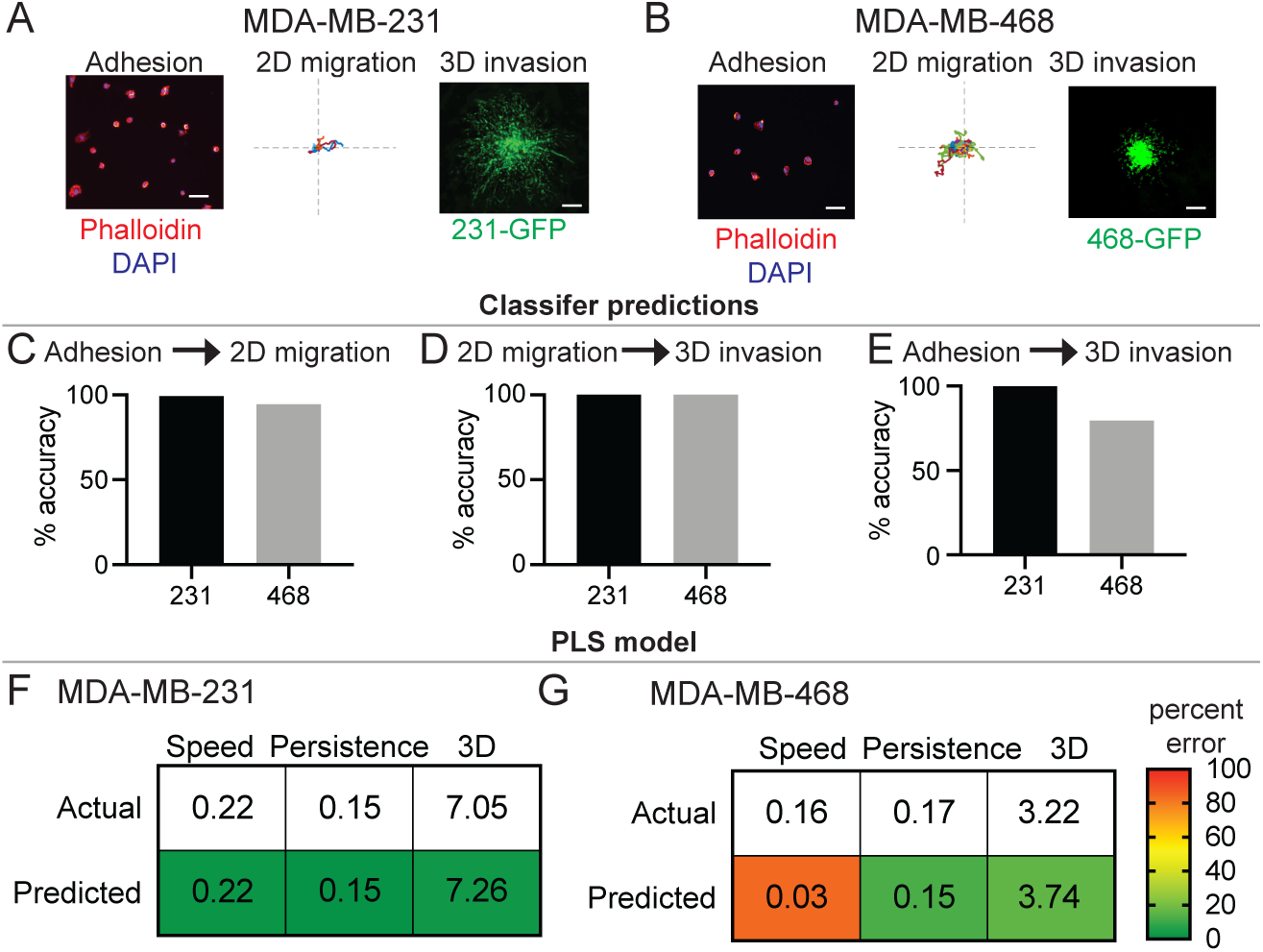
Classifier and PLS models accurately predict response to Matrigel in breast cancer cells. Representative cell adhesion, 2D migration, and 3D invasion of MDA-MB-231 (A) and MDA-MB-468 (B) cells on or in Matrigel. Representative cell adhesion images show cells fixed and stained with Phalloidin (red) and DAPI (blue) after 2hrs on Matrigel. Scale bar is 100μm. 2D migration roseplots show individual cell tracks on Matrigel for 16hrs. Axis length is 500 μm. Representative 3D invasion spheroids are made from 231-GFP and 468-GFP cells embedded in Matrigel gels for 5 days. Scale bar is 200μm. Accuracy of classifier models to predict 2D cell migration of cells on Matrigel from all cell adhesion parameters (C) and 3D invasion of cells in Matrigel from 2D cell migration speed and persistence (D) and all cell adhesion parameters (E). Best classifier model was used to predict. Linear SVM for C, Linear SVM and SVM for D, and Linear SVM and RF for E. Prediction of 2D cell migration speed, 2D persistence, and 3D invasion of cells on Matrigel from MDA-MB-231 (F) and MDA-MB-468 (G) PLS models built with 2 principal components. Numbers represent the actual and predicted values for each metric. Colors represent percent error, where green is a low error and red is a high error, indicated by the color gradient.

We next tested the ability of our PLS models to predict the responses of new cell lines. For these experiments, we used BT-549, another human triple-negative breast cancer cell line. We evaluated the effects of Fibronectin, Tenascin C, and Matrigel on cell adhesion, 2D migration, and 3D invasion of BT-549 cells (Fig 8A-D). We find that these ECM proteins have distinct effects on the shape, migration, and invasion of the cells, consistent with our previous data. Next, we evaluated how well the MDA-MB-231, MDA-MB-468, and combined models predict the BT-549 responses to these ECM proteins. We did this with and without training the models with BT-549 data (Fig 8E). We find that when we trained the original models with BT-549 response to no ECM, the error reduced. Training the models with BT-549 responses to both no ECM and Fibronectin also increases how accurately Tenascin C and Matrigel are predicted, as seen by the lower percent error (Fig 8E). Interestingly, the MDA-MB-231 model predicts BT-549 response to Tenascin C and Matrigel better than the MDA-MB-468 model. We also find that the models more accurately predicts responses to Tenascin C than Matrigel.

**Figure 8:**
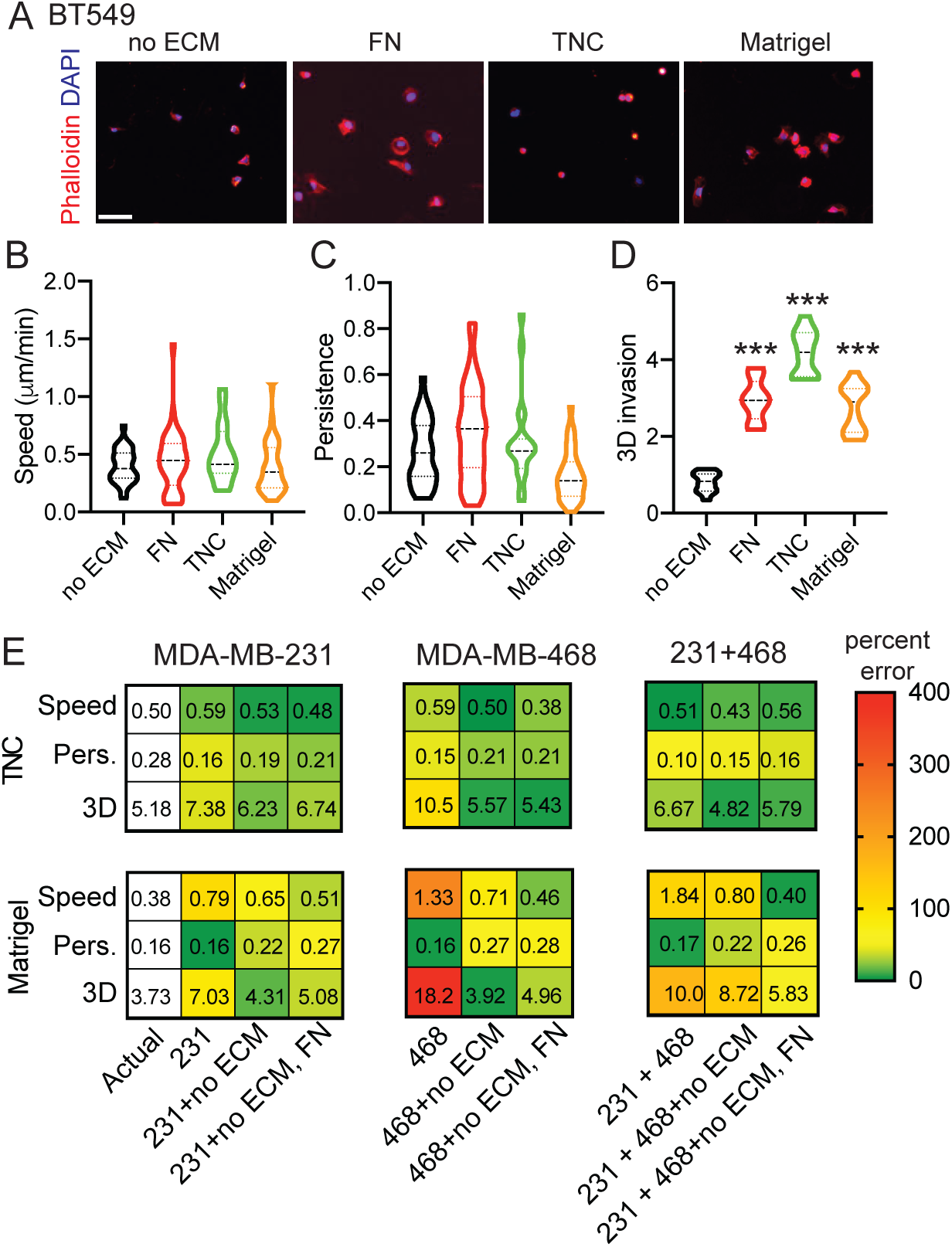
ECM-driven predictions are cell-line specific. Cell adhesion (A), 2D migration (B,C), and 3D invasion (D) of BT-549 cells on three ECM substrates: Fibronectin, Tenascin C, and Matrigel. A) Representative cell adhesion images show cells fixed and stained with Phalloidin (red) and DAPI (blue) after 2hrs on Matrigel. Scale bar is 100μm. Quantification of cell migration speed (μm/min) (B) and persistence (C). D) Quantification of fold change in BT-549-GFP spheroid area on day 5 relative to day 1. Data pooled from 1 biological replicate, with technical triplicates. *** p<0.001 by one-way ANOVA and Dunn’s multiple comparison test. E) Prediction of 2D cell migration speed, 2D persistence, and 3D invasion of BT-549 cells on Tenascin C and Matrigel from MDA-MB-231, MDA-MB-468, and combined PLS models built with 2 principal components for one cell line and 6 principal components for combined cell lines. Predictions were done without BT-549 data, with the addition of BT-549 no ECM, and with the addition of BT-549 no ECM and Fibronectin. Numbers represent the actual and predicted values for each metric. Colors represent percent error, where green is a low error and red is a high error value, indicated by the color gradient.

## Discussion

Our goal was to identify the relationship between ECM responses in adhesion, 2D migration and 3D invasion assays to develop strategies to easily predict the effect of novel ECM proteins on 3D cancer cell invasion, which is more relevant to the study of cancer metastasis. 3D invasion assays can be more complex, time consuming and more challenging for follow up analysis, while 2D migration and adhesion assays are quicker, easy to analyze, and to use with other experimental approaches such as cell sorting, atomic force microscopy or immunostaining. By evaluating the response of two TNBC breast cancer cell lines to 4 ECM proteins known to be upregulated in metastatic breast cancers, we found that there is no linear relationship between metrics used to quantify these 3 assays. Using machine learning algorithms, we found that cell adhesion can successfully and accurately classify 2D migration speed and 3D invasion, while 2D migration speed and persistence are unable to classify ECM-driven 3D invasion. ECM proteins have distinct effects on cell adhesion, which is characterized by features that characterize cell size, irregularity and elongation. Using data-driven modeling, we find that some shape parameters, such as those that quantify cell elongation and irregularity, are more important for predicting 2D migration and 3D invasion. Finally, we show that both methods can be used to accurately predict the effect in 2D and 3D of a new ECM protein, but that these predictions are cell-line specific and that models generated for one cell line cannot be used for another. Overall, these studies suggest that the shape a cell takes in response to an ECM protein, and not 2D migration speed, is more predictive of 3D invasion and our data provides a pipeline to predict the effect of novel ECM proteins in driving invasion and metastasis based on a simple adhesion assay.

In this study, we found that both machine learning classifier models and data-driven PLS models could be used to accurately predict 2D and 3D responses of cells to a new ECM protein. However, we found that classifier models did not classify all cells of the new ECM protein data set accurately and the results varied per classifier algorithm used (Fig S8 A-F). Additionally, the algorithms that scored the highest ROC scores during training and validation would not necessarily predict with highest accuracy when tested. RF and LR algorithms performed the weakest for ECM prediction, which could be due to issues with overfitting or suitability for complex relationships with multiple parameters, respectively. We find that SVM algorithms worked best here, which can model non-linear decision boundaries, are robust against overfitting, and work well for smaller datasets like ours. The limitations of predictions with classifier models could be addressed by using a more quantitative PLS model. We find that our PLS model was able to quantitatively define the relationships between cell adhesion and the 2D and 3D responses of cells by iteratively reducing the dimensionality of the training data set. With PLS modeling, we are able to extract which cell shape parameters are most strongly connected with cell responses in 2D and 3D, which allows us to generate hypotheses and quantitatively support them (17). Nevertheless, the PLS model still had important limitations. When predicting the effects of Tenascin C and Matrigel in the new cell line, the PLS model was more accurate at predicting the effects of Tenascin C. Matrigel is known to be a mixture of several growth factors and ECM proteins, suggesting that its effect on cell migration is more complex. Indeed, we have shown that there can be synergy between growth factors and ECM protein, suggesting that combinations of cues from multiple growth factors and ECM proteins can lead to more complex results (26). We also found that the PLS model constructed with MDA-MB-231 cells predicted ECM-driven effects in BT-549 cells better than the PLS model with MDA-MB-468 cells. MDA-MB-231 cells are known to be more mesenchymal, and it has been found that BT-549 cells express characteristics of more mesenchymal cells, as seen by lower surface levels of Integrin B4 (24). BT-549 was also found to have a similar metastatic potential to MDA-MB-231 cells, suggesting that the two mesenchymal cell lines would respond similarly to different ECM proteins (25). Therefore, our model may be best suited for studies within a single cell line and with individual ECM proteins. Future studies will have to address how combinations of ECM cues, with and without other pro-migratory such as growth factors, impact cancer cell invasion.

These studies also shed light on the heterogeneity of responses to ECM proteins in all these assays, particular for the adhesion and migration, where the data is quantified at the single cell level. Some ECM proteins, like Tenascin C and Collagen I, induce more homogeneous responses in terms of cell shape parameters, while others like Fibronectin and Collagen IV have a range of effect (Fig 2). Previous studies have linked heterogeneity of cell shape to metastatic potential. For example, lower variation in morphology is predictive of cells derived from metastatic sites, but not associated with any particular somatic mutations (27). In addition, the dynamics of breast cancer cell shape heterogeneity can impact response to therapy. Indeed, time series modeling that captures the heterogeneous dynamic cellular responses can improve drug classification and provide insight into mechanisms of drug action (28). The mechanisms that govern this heterogeneity in breast cancer remain poorly understood. We have shown that changes in alternative splicing of the actin regulator Mena can impact sensitivity to FN gradients *in vivo*. Breast cancer cells that express the Mena^INV^ isoform, which includes a 19 amino acid exon, are more sensitive to FN which increases their metastatic potential (5, 29). Expression of Mena^INV^ is regulated by the acidity of local environment (30), suggesting that feedback between the tumor microenvironment and the cancer cells themselves is critical in regulating the signaling pathways that will impact cell shape heterogeneity. It will be important to evaluate how the heterogeneity of response to ECM proteins impacts metastasis and response to therapy.

Migration responses in 2D are quantified with two main metrics: cell migration speed and persistence, however, it is not clear what migration response is more relevant to metastasis. Interestingly, in the World Cell Race, speed and persistence correlated for the migration of over 50 cell types on fibronectin coated lines (31). In a follow-up study, persistence was found to be robustly coupled to cell migration speed (32), however these studies were done in epithelial and myeloid cells and are not in response to a given cue. We find no correlation between ECM-driven cell speed and persistence, and that cell shape is more predictive for persistence than it is of cell speed. Persistence may be more relevant to study in response to a directional cues, such as in the context of haptotaxis or chemotaxis (3, 33). For example, directed migration of breast cancer cells to gradients of Fibronectin increases directional persistence to promote metastasis, without affecting cell speed (5). Therefore, the nature and organization of the cue driving cell migration may play an important role in determining which metric is more predictive of metastasis potential. Our studies further demonstrate that in the context of ECM responses, cell shape is predictive of 3D invasion, with ECM-driven effects on 2D speed not predictive of 3D invasion. Interestingly, these data are similar to what was found for response to growth factors, another pro-migratory cue. Whereas 2D migration properties did not correlate well with 3D behavior across multiple growth factors, Meyer *et al.* found that increased membrane protrusion elicited by growth factor stimulation did relate robustly to enhanced 3D migration properties in several breast cancer cells (34). These studies further support the importance of considering the properties of the cue to best evaluate its role on breast cancer metastasis. Future studies should address the importance of cell speed, persistence and invasion to metastasis *in vivo*.

## Supplemental Figures

**Figure S1:**
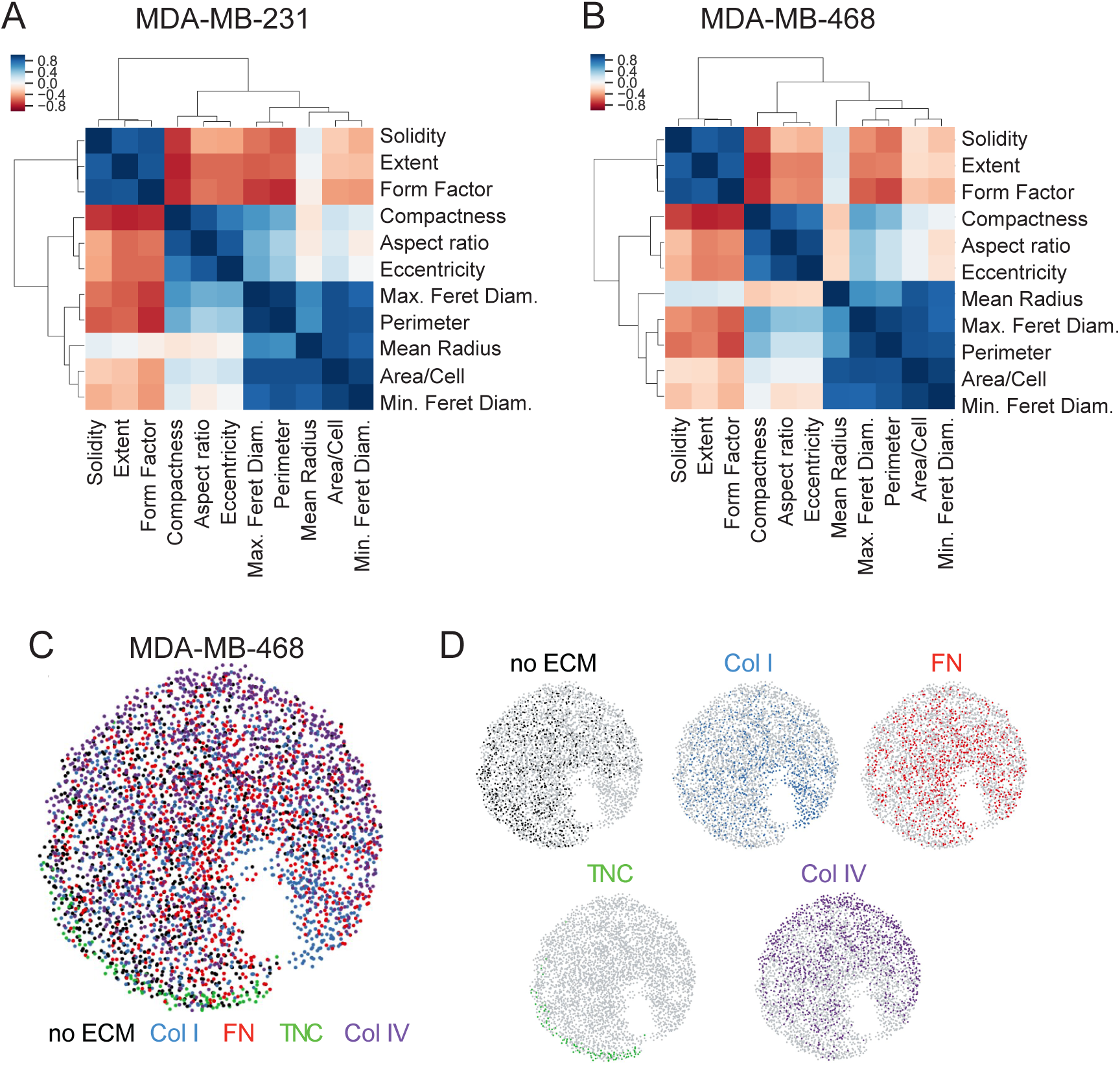
Analysis of cell shape parameters. Correlation between 11 shape parameters for MDA-MB-231 (A) and MDA-MB-468 (B) cells. Each cell adhesion parameter is clustered by rank correlation and mean linkage. C) Visualization of continuum of cell adhesion of MDA-MB-468 cells on different ECM substrates based on all 11 cell shape parameters with SPRING plots. D) Plots showing localization of cell adhesion shape on each ECM factor on the combined SPRING plot.

**Figure S2:**
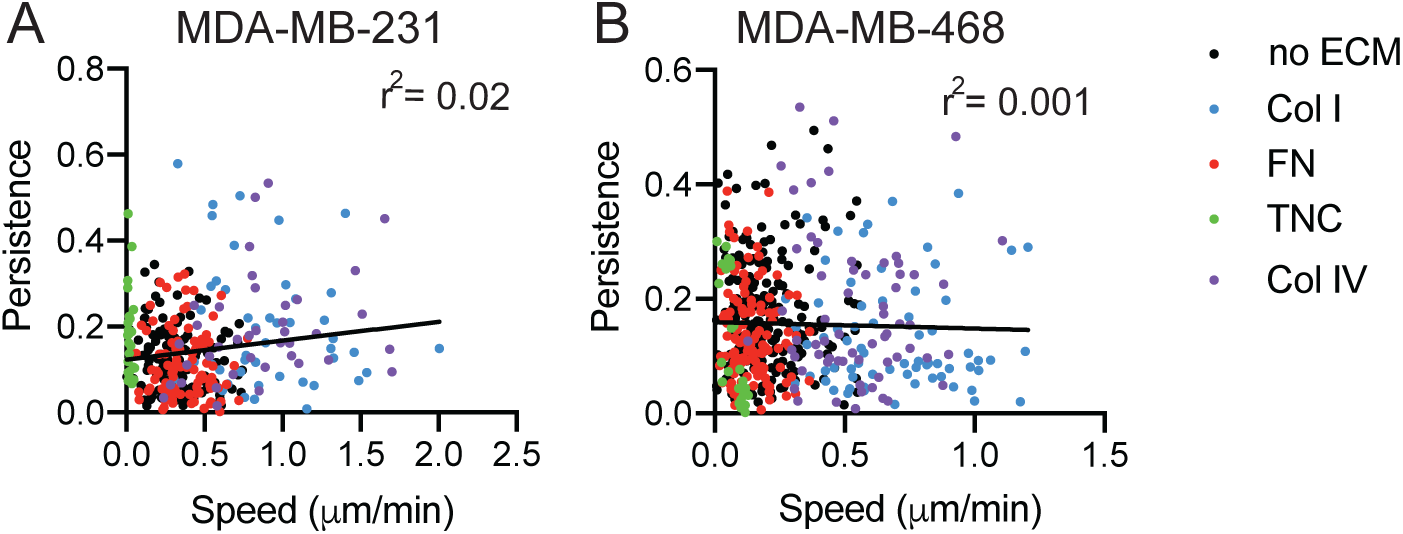
ECM-driven effects on 2D cell migration speed and 2D persistence do not correlate for individual cells. Correlation between 2D cell migration speed and 2D persistence of each cell tracked for MDA-MB-231 (A) and MDA-MB-468 (B) cells. Data pooled from at least 3 independent experiments, between 30-130 individual cells analyzed per condition.

**Figure S3:**
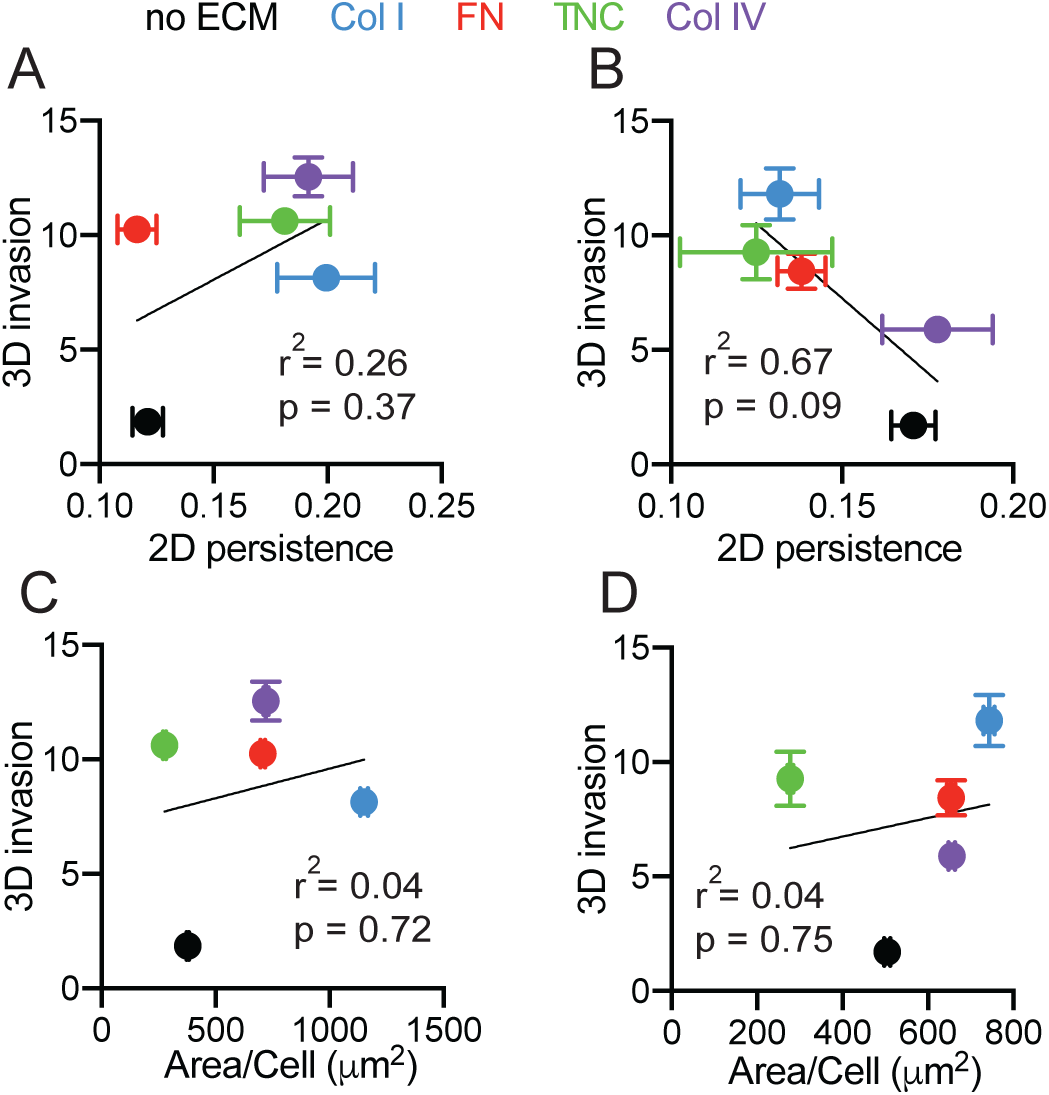
ECM-driven 3D invasion does not correlate with persistence or cell shape. Correlation between mean fold change in spheroid area and 2D persistence for 231-GFP (A) and 468-GFP (B) cells embedded in media, Collagen I, Fibronectin, Tenascin C and Collagen IV. Correlation between mean fold change in spheroid area and cell area for 231-GFP (C) and 468-GFP (D) cells embedded in media, Collagen I, Fibronectin, Tenascin C and Collagen IV.

**Figure S4:**
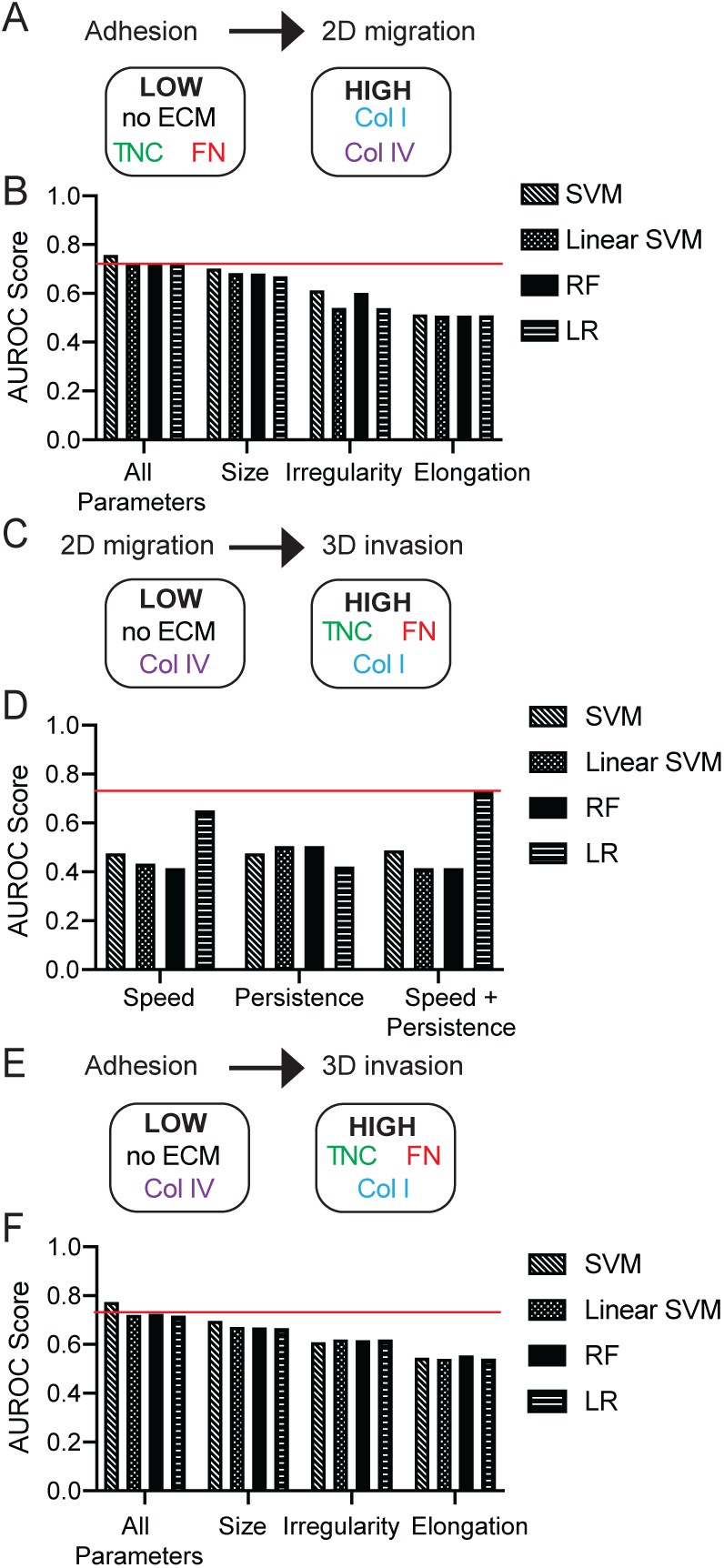
Adhesion classifies ECM-driven 2D migration and 3D invasion in MDA-MB-468 cells. A) Cell adhesion to predict binary classification of 2D cell migration speed of MDA-MB-468 cells on plastic, Collagen I, Fibronectin, Tenascin C, or Collagen IV. B) AUROC scores of binary classifier models (A) using all 11 cell shape parameters, cell size parameters (area/cell, perimeter, mean radius, min feret diameter, max feret diameter), cell irregularity parameters (solidity, extent, form factor), and cell elongation parameters (eccentricity, aspect ratio, compactness). C) 2D cell migration to predict binary classification of mean fold change of spheroid area of 468-GFP cells embedded in media, Collagen I, Fibronectin, Tenascin C, or Collagen IV. D) AUROC scores of binary classifier models (C) using 2D cell migration (cell migration speed and persistence). E) Cell adhesion to predict binary classification of mean fold change of spheroid area of 468-GFP cells embedded in media, Collagen I, Fibronectin, Tenascin C, or Collagen IV. F) AUROC scores of binary classifier models (E) using all 11 cell shape parameters, cell size parameters, cell irregularity parameters, and cell elongation parameters.

**Figure S5:**
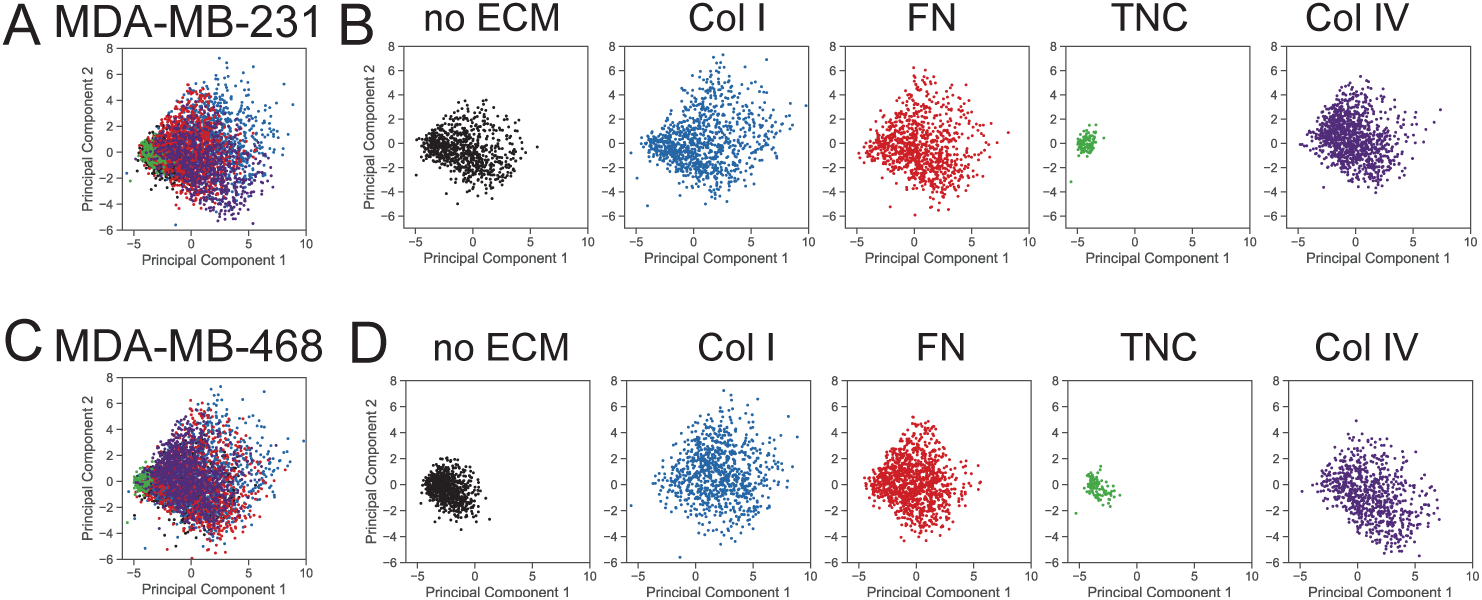
PCA analysis for MDA-MB-231 and MDA-MB-468 cells. A) Principal component analysis (PCA) of covariance between different metrics to quantify MDA-MB-231 adhesion on plastic, Collagen I, Fibronectin, Tenascin C or Collagen IV, indicated by the colors in the legend. B) PCA plot showing localization of each ECM substrate on same axes as A. C) PCA of covariance between cell adhesion metrics of MDA-MB-468 cells on plastic, Collagen I, Fibronectin, Tenascin C or Collagen IV. D) PCA plot showing localization of each ECM substrate on same axes as C.

**Figure S6:**
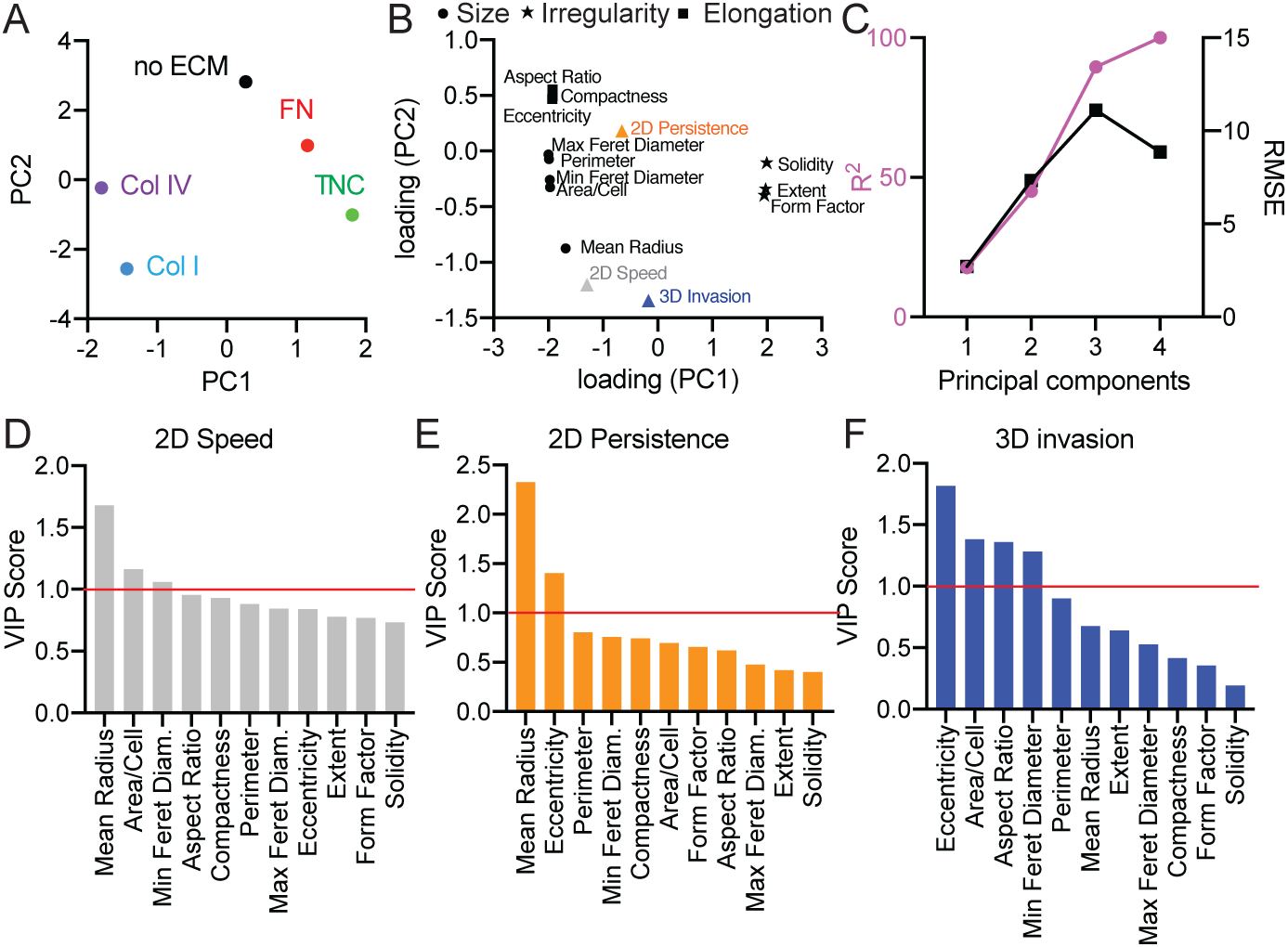
PLS model constructed to predict ECM-driven 2D migration and 3D invasion from cell adhesion of MDA-MB-468 cells. A) Scores plot for PLS model with MDA-MB-468 cells. Principal components reflect covariation between adhesion, 2D migration and 3D invasion for each cell line on each ECM protein. B) PLS loading plot for 11 cell adhesion shape parameters and 3 cell responses. C) R^2^ and root mean square error (RMSE) for the PLS model built with increasing numbers of principal components. Ranked VIP scores for predicting 2D cell migration speed (D), 2D persistence (E), and 3D invasion (F). VIP score >1 indicates important cell shape parameters to predict cell invasion response.

**Figure S7:**
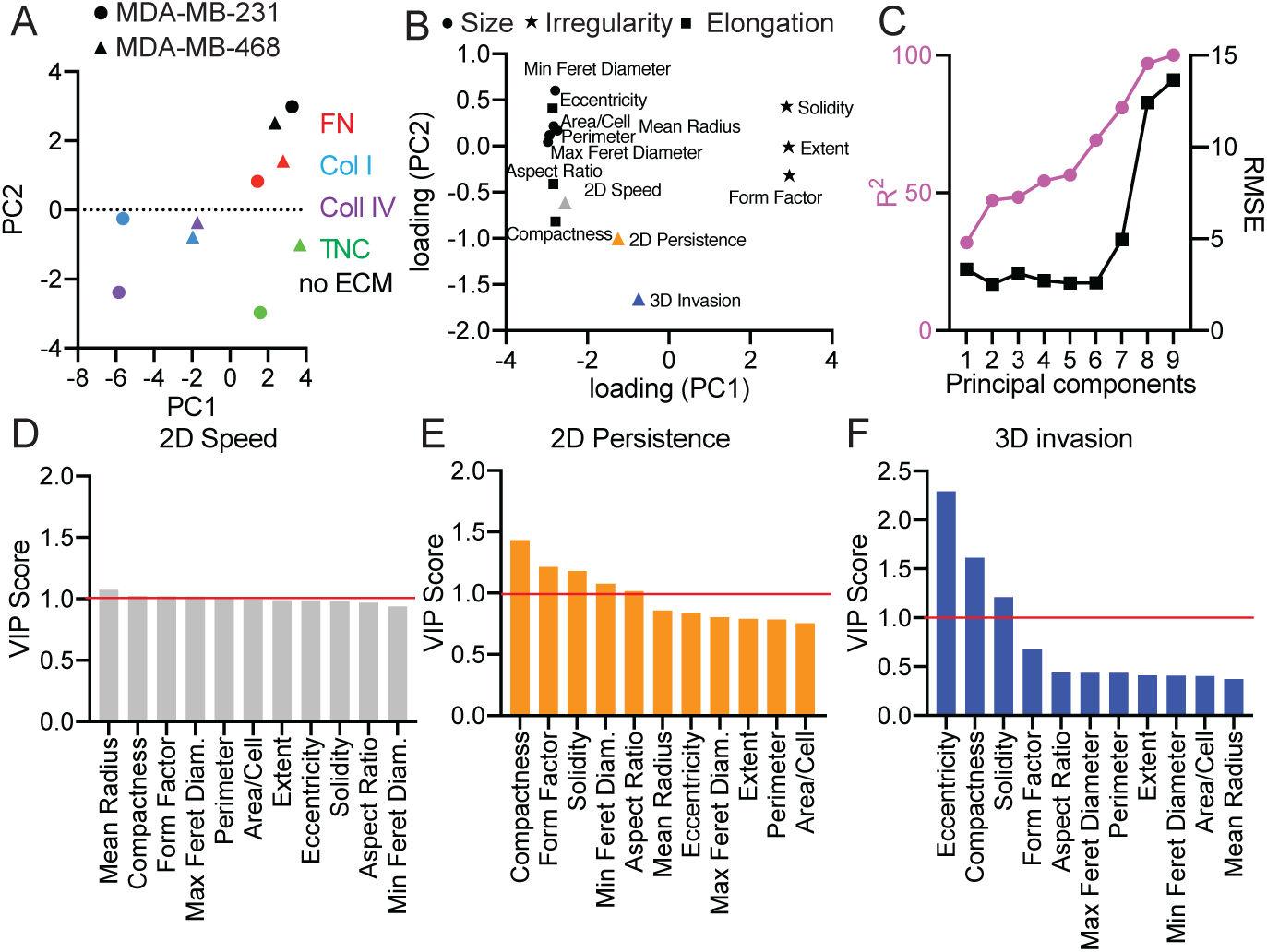
PLS model constructed to predict ECM-driven 2D migration and 3D invasion from cell adhesion of both MDA-MB-231 and MDA-MB-468 cells. A) Scores plot for PLS model with both MDA-MB-231 and MDA-MB-468 cells. Principal components reflect covariation between adhesion, 2D migration and 3D invasion for each cell line on each ECM protein. B) PLS loading plot for 11 cell adhesion shape parameters and 3 cell responses. C) R^2^ and root mean square error (RMSE) for the PLS model built with increasing numbers of principal components. Ranked VIP scores for predicting 2D cell migration speed (D), 2D persistence (E), and 3D invasion (F). VIP score >1 indicates important cell shape parameters to predict cell invasion response.

**Figure S8:**
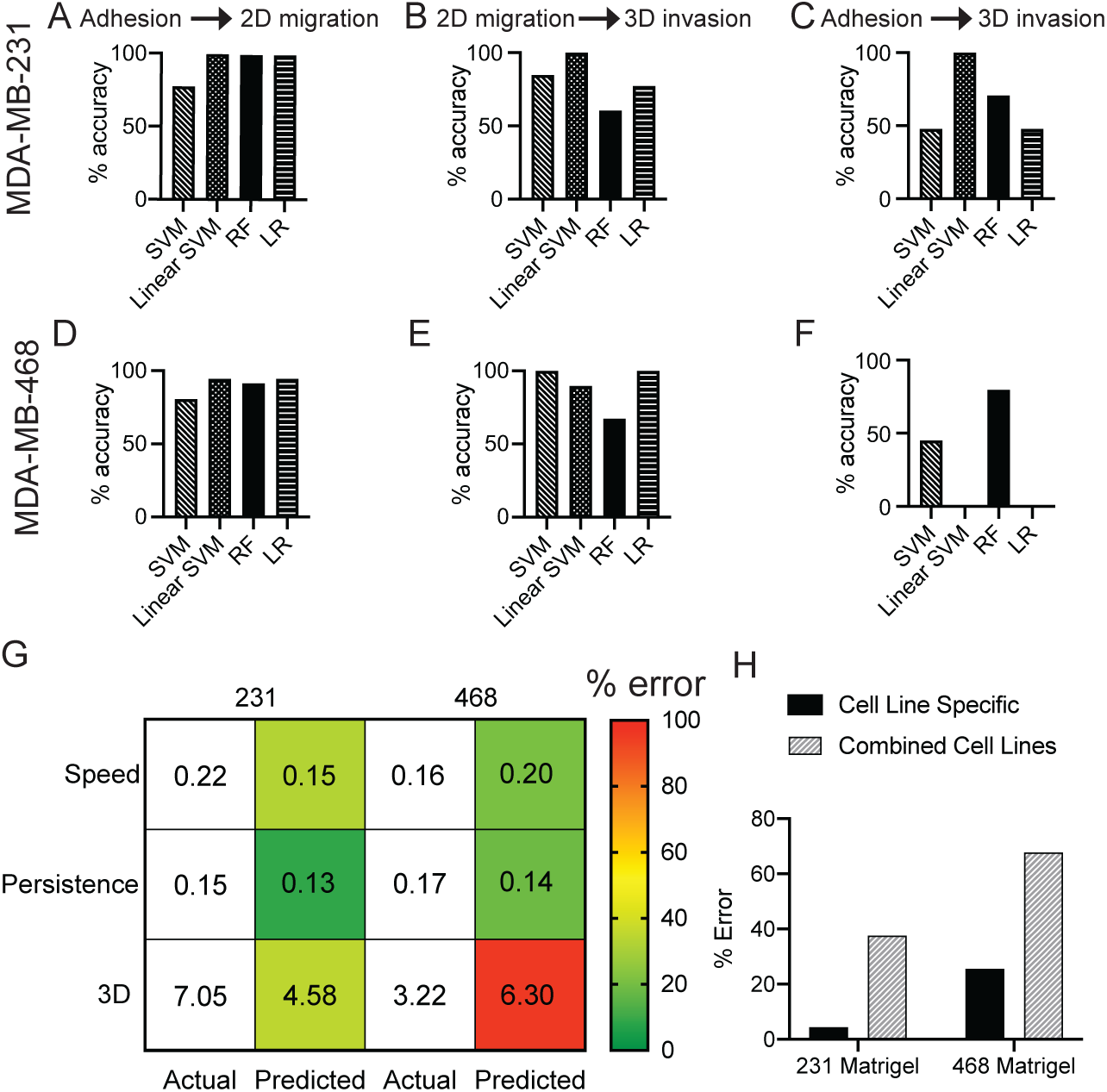
Classifier and PLS predictions for Matrigel-driven responses. Accuracy of prediction of 2D migration (A, D) and 3D invasion (B, C, E, F) of MDA-MB-231 (A-C) and MDA-MB-468 (D-F) cells in Matrigel using binary classifier models. G) Prediction of 2D cell migration speed, 2D persistence, and 3D invasion of cells on Matrigel from cell adhesion using combined cell line PLS model built with 6 principal components. Numbers represent the actual and predicted values for each metric. Colors represent percent error, indicated by the color gradient. H) Percent error of 3D invasion prediction of 231-GFP and 468-GFP cells in Matrigel from both cell adhesion and 2D migration. Predictions are made within the same cell line and with the combined cell line PLS model.

